# Robust memory of face moral values is encoded in the human caudate tail: A simultaneous EEG-fMRI study

**DOI:** 10.1101/2023.07.22.550131

**Authors:** Ali Ataei, Arash Amini, Ali Ghazizadeh

## Abstract

Moral judgements about people based on their actions is a key component that guides social decision making. It is currently unknown how positive or negative moral judgments associated with a person’s face are processed and stored in the brain. Here, we investigate the long-term memory of moral values associated with human faces using simultaneous EEG-fMRI data acquisition. Results show that only a few exposures to morally charged stories of people, are enough to form long-term memories a day later for a relatively large number of new faces. Event related potentials (ERPs) showed a significant differentiation of remembered good vs bad faces over centerofrontal electrode sites (value ERP). EEG-informed fMRI analysis revealed a subcortical cluster centered on the left caudate tail (CDt) as a correlate of the face value ERP. Importantly neither this analysis nor a conventional whole brain analysis revealed any significant activation in cortical areas in particular the fusiform face area (FFA). Conversely an fMRI-informed EEG source localization using accurate subject-specific EEG head models also revealed activation in the caudate tail. Nevertheless, the detected caudate tail region was found to be functionally connected to the FFA, suggesting FFA to be the source of face-specific information to CDt. These results identify CDt as the main site for encoding the long-term value memories of faces in humans suggesting that moral value of faces activates the same subcortical basal ganglia circuitry involved in processing reward value memory for objects in primates.

## Introduction

Our past experiences with people whether good or bad affect our future interactions with them and guide our social decision making. In particular, moral judgement about people based on their actions is a key component in our evaluation of an individual (Cornwell & Higgins, 2019; Jiang et al., 2022). It is currently unknown how positive or negative moral judgments associated with a person’s face are processed and stored in the brain in long-term memory.

Human brain has specialized face processing areas in the temporal cortex including the occipital face area (OFA), superior temporal sulcus (STS) and the fusiform face area (FFA) (Kanwisher & Yovel, 2006). Several studies have examined the coding of various aspects of faces based on their intrinsic visual features such as emotional expressions (Engell & Haxby, 2007; Winston et al., 2004), attractiveness (Cloutier et al., 2008; Kranz & Ishai, 2006; O’Doherty et al., 2003; Said et al., 2011; Winston et al., 2007) trustworthiness (Engell et al., 2007; Winston et al., 2002) or social value (Oosterhof & Todorov, 2008; Todorov et al., 2011) in the human brain with some divergent results. A recent review and meta-analysis of these findings suggests consistent activations for negative evaluations in the amygdala, and for positive evaluations in the medial orbitofrontal cortex (mOFC), anterior cingulate cortex (ACC), caudate nucleus and the nucleus accumbens (NAcc) (Mende-Siedlecki et al., 2013). None of these studies however, address evaluations based on explicit value association with faces in long-term memory and independent of visual features.

The value circuitry for objects in general is extensively studied in both humans and non-human primates with key cortical and subcortical areas such as orbitofrontal cortex (OFC), insula, ACC, basal ganglia, amygdala and midbrain dopaminergic areas being activated during object reward association tasks (Berridge & Kringelbach, 2008; Kim et al., 2014; Morrison & Salzman, 2010; Padoa-Schioppa & Assad, 2006; Rushworth & Behrens, 2008; Schultz, 2007). Value memory is also shown to activate a temporal-prefrontal circuitry along with its functionally connected subcortical areas, in particular, the caudate nucleus, amygdala and claustrum (Ghazizadeh, Griggs, et al., 2018; Ghazizadeh, Hong, et al., 2018; Ghazizadeh & Hikosaka, 2021; Kang et al., 2021; Kim & Hikosaka, 2013; Yasuda et al., 2012). It is not known how long-term memory of associated value with faces engages this circuitry and whether it can change even the primary face processing in areas such as FFA.

To address this question, we randomly assigned positive or negative values based on morally charged biographical stories to novel faces and then examined the brain activations to these faces a day later using a simultaneous EEG-fMRI paradigm which involved a binary choice for face values. The value ERP showed a significant differentiation of the truly identified good and bad faces over center-frontal electrodes, peaking at about 0.6s post-stimulus and lasting until the end of face presentation. This ERP is shown to originate from the left caudate tail, based on results from both EEG-informed fMRI analysis and fMRI-informed EEG source localization. Interestingly, while none of the cortical face processing areas were found to encode the face values, some were found to be functionally connected to the detected value-coding caudate tail region.

## Results

To create value memory for faces, we used 24 arbitrary artificial faces created by StyleGAN2 (Karras et al., 2019). Each face was randomly assigned to a brief unique biography with either a positive (good faces) or a negative (bad faces) moral value (see supplementary table 1 for a list of all stories). In the value training session, subjects viewed each face while listening to its short story. Each face was viewed for 10 sec within a block of 24 faces and the process was repeated 2 times, 2 hours apart. The assignment of biographies to the faces were swapped across so that each face was associated with positive values for half of the subjects and with negative values for the other half.

One day later, we tested the face value memory of subjects in the MRI scanner being simultaneously equipped with an MRI-compatible EEG cap (value memory session). During the experimental task, each face was portrayed for 2.5s (Fig. 1b). Then, a black page was shown with two letters of “G” and “B” (referring to “Good” and “Bad”, respectively) on the left or right visual hemifield for 2s, during which, the subject had to indicate his/her response by pressing a response button using the corresponding hand. The location of letters “G” and “B” were randomly flipped in each trial. The subjects were instructed to answer all trials based on their closest guess, even if they did not remember an explicit history about a face. The subjects’ accuracy in identifying face types was above 72% which is significantly higher than the chance level for both good and bad faces (Fig. 1c; t-test, p < 1e-6 & p < 1e-8, for bad and good faces respectively). Nevertheless, the performance for the good faces was significantly better than that for bad ones by about 12% (Fig. 1c; paired t-test, p = 0.004). This suggests that a brief exposure to a large number of new people (24 new faces) and their moral stories, creates a lasting memory that is accessible a day later.

**Figure 1:**
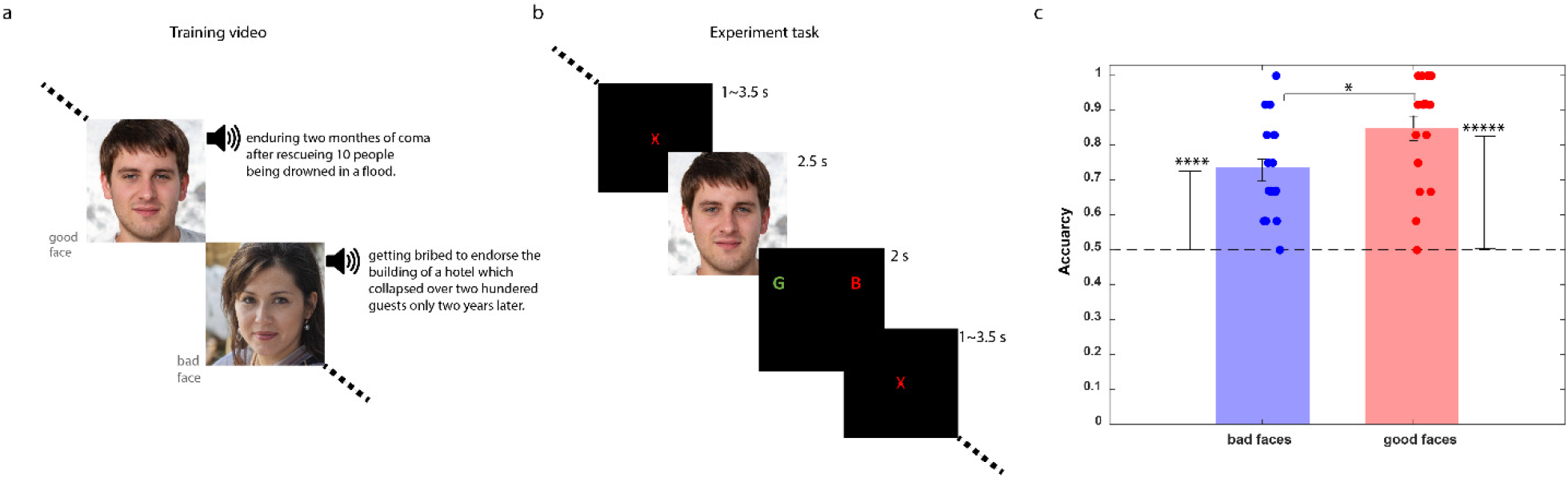
Subjects’ training procedure, memory task and behavioral results. **a**) Value training: the subjects watched a 5-minute video clip two times, the night before the experiment. In this video, 24 faces (produced by a deep generative network) were associated with short biographical stories. Each story contained a positive or negative moral value about the presented face (good or bad faces, see suppl. Table 1 for the list of morally-charged stories). **b**) Memory task: faces seen the day before were shown in a random sequence for each subject. Each face was shown for 2.5s. Then, a black page was shown with two letters of “G” and “B” (referring to “Good” and “bad”, respectively) for 2s, during which, the subject had to indicate his/her response by pressing the appropriate button. The sides of letters “G” and “B” were randomly filliped in each trial. Then, an inter-stimulus-interval (a black page with a red fixation cross at center) followed for a random time period between 1 to 3.5s. **c)** Behavioral result: Performance of subjects in judging the value of good and bad faces, along with the chance level (=50%, dashed line) is shown. The subjects’ performance for both categories were significantly higher than chance level (t-test, p < 1e-6 & p <1e-8 for bad and good faces, respectively) while they had remembered good faces 12% better than bad ones on average (t-test, p < 0.004). Individual subject data points are shown. Error bars indicate standard errors.

### Robust differentiation of remembered good and bad faces over center-frontal electrodes

To study the differential neuronal activity during remembered good and bad faces, event-related potentials (ERPs) for each category, separately were calculated. In order to create a more precise group average, the EEG signals for each subject were normalized and transferred to the standard cap situated over the standard brain (Fig. 2a, see methods). The ERPs for the remembered good and bad faces (value ERP) showed a robust differentiation between them over a large portion of the 2.5s stimulus presentation on center-frontal channels such as C1 and FC1 (t-test against baseline; p-value ≤0.001, Fig. 2a). To ensure quality of EEG responses, group-average of the visual ERPs was also calculated which showed a robust signal over the occipital electrodes (suppl. Fig. 1).

**Table 1:** activated clusters. Sizes, coordinates and cluster-correction thresholds for the activated clusters.

|  | cluster size<br>(voxels) | CM x | CM y | CM z | peak x | peak y | peak z | cluster threshold<br>for p=0.01<br>(voxels) |
| --- | --- | --- | --- | --- | --- | --- | --- | --- |
| corr. good -corr. Bad: |  |  |  |  |  |  |  | 66 |
| L caudate tail | 86 | 29.3 | 38.3 | -1.2 | 32 | 38 | -4 |  |
| EEG-driven regressor |  |  |  |  |  |  |  | 51 |
| L caudate tail | 56 | 31.7 | 28.2 | -8.2 | 34 | 26 | -8 |  |
| correct - incorrect: |  |  |  |  |  |  |  | 74 |
| L angular gyrus | 296 | 47.7 | 59.5 | 35.1 | 46 | 56 | 34 |  |
| L superior frontal gyrus | 151 | 18.7 | -27.8 | 52.2 | 18 | -28 | 58 |  |
| L medial temporal gyrus | 75 | 64 | 25.9 | -7.3 | 62 | 26 | -8 |  |
| right hand - left hand |  |  |  |  |  |  |  | 80 |
| R motor cortex | 1393 | -37.8 | 25 | 56.1 | -52 | 16 | 56 |  |
| L cerebellum | 655 | 17.5 | 52.7 | -21 | 26 | 50 | -22 |  |
| L lingual cortex | 583 | 19.2 | 65 | -8.9 | 16 | 74 | -10 |  |
| R insula | 447 | -45.6 | 16.4 | 13.9 | -48 | 14 | 16 |  |
| R cerebellum | 328 | -20.6 | 48.9 | -22.5 | -20 | 50 | -22 |  |
| L motor cortex | 318 | 36.9 | 23.3 | 57.5 | 36 | 22 | 44 |  |
| L cerebellum2 | 261 | 16.3 | 60.3 | -47 | 16 | 58 | -46 |  |
| R cingulate gyrus | 254 | -7.6 | 10.4 | 50 | -10 | 26 | 48 |  |
| R posterior cingulate | 242 | -12.7 | 59.8 | 3.6 | -4 | 74 | 8 |  |
| R Cuneus | 215 | -13.1 | 93.7 | 4 | -16 | 94 | 0 |  |
| L Cuneus | 200 | 19 | 86.3 | 23.9 | 12 | 86 | 28 |  |
| L Cuneus2 | 164 | 8 | 76.8 | 9.5 | 10 | 70 | 6 |  |
| R thalamus | 149 | -10.9 | 18.4 | 9.7 | -10 | 20 | 12 |  |
| R superior temporal gyrus | 117 | -55.5 | 16.1 | 7.6 | -50 | 18 | 10 |  |
| L Cuneus3 | 112 | 3 | 87.4 | 10.5 | 0 | 90 | 18 |  |
| R inferior frontal gyrus | 84 | -59.7 | -7.3 | 17.9 | -60 | -12 | 22 |  |

**Figure 2:**
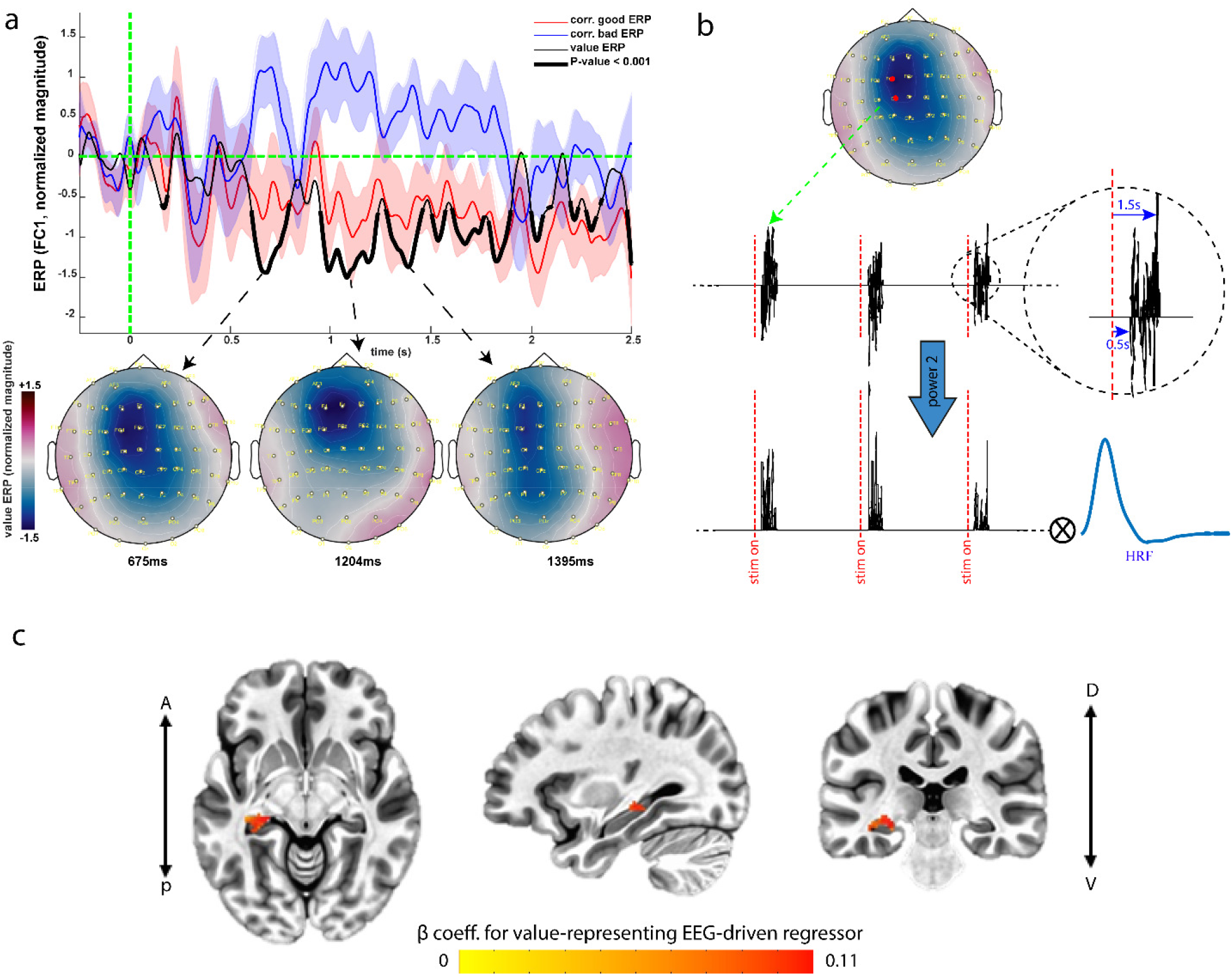
Value ERP & the EEG-informed fMRI analysis. **a)** The group-average ERPs for the memorized good and bad faces and their difference (value ERP) in electrode FC1 along with the associated group-average scalp potential topographies at some sample time points. Parts of the value ERP that are significantly different from its baseline are shown with bold line (p-value < 0.001, uncorrected), **b)** The average of electric potentials in center-frontal electrodes over the significant interval (0.5-1.5s post-stimulus) is considered as the EEG-driven value signal. This signal is then squared and consequently convolved with the canonical HRF to build our EEG-driven regressor. **c)** EEG-informed fMRI analysis: the group-average of the beta coefficients for the EEG-driven regressor shows significant activation in a cluster centered on the left caudate tail (p-value < 0.01, cluster-corrected).

### EEG-informed fMRI analysis reveals caudate tail as the origin of the value ERP

We next performed an EEG-informed fMRI analysis. Based on the time course of value ERP, we used the average EEG signal over C1 and FC1 in the interval from 0.5s to 1.5s post-stimulus as the EEG regressor in the fMRI GLM model (Fig.2b, see methods). We have previously shown that since the BOLD signal measures the energy consumption in the brain, the square (or power 2) of the EEG signals is the optimal regressor of the BOLD responses (Ataei et al., 2022). Thus, to localize the source(s) of EEG signals in the center-frontal electrodes, its square was used as the *trial by trial* correlate of face value signal in the brain (Fig.2b).

Other regressors in the model included correct answer to good faces, “correct good”, correct answer to bad faces, “correct bad”, incorrect answer to good faces, “incorrect good” and incorrect answer to bad faces, “incorrect bad”. In case of no incorrect answer to a certain category, the associated regressor was omitted for that subject. Moreover, to account for the effect of subject’s motor action during his/her button press response, we also added two confounding regressors, one for each hand. Finally, the average BOLD signal inside the ventricles was used as another nuisance regressor to ensure that responses in structures near the ventricles are not affected by such nonneural extraneous signals (GLM1, see materials & methods for details of the regressor design). As a control analysis, the right versus left hand contrast showed significant activation in the contralateral and deactivation in the ipsilateral motor cortices (suppl. Fig. 2a, p-value < 0.01, cluster-corrected). Moreover, activation of the ipsilateral and deactivation of the contralateral cerebellar cortices were also observable (suppl. Fig. 2b, p-value < 0.01, cluster-corrected), consistent with existing literature (Thickbroom et al., 2003).

The BOLD correlate of center-frontal EEG regressor showed significant activation only in a subcortical cluster centered on the left caudate tail (CDt) (Fig. 2c, p-value < 0.01, cluster-corrected). Notably, the contrast of correct good v.s. correct bad (the value contrast) did not reveal any significant activation in the brain suggesting that the source of observed EEG responses in the center-frontal electrodes were most likely limited to CDt and that this activity was well captured by the EEG power of center-frontal electrodes in each trial. This is expected since a correct model of statistical dependence among variables predicts that given the EEG signal originating from value contrast, BOLD should become independent of value contrast itself (suppl. Fig. 3a).

**Figure 3:**
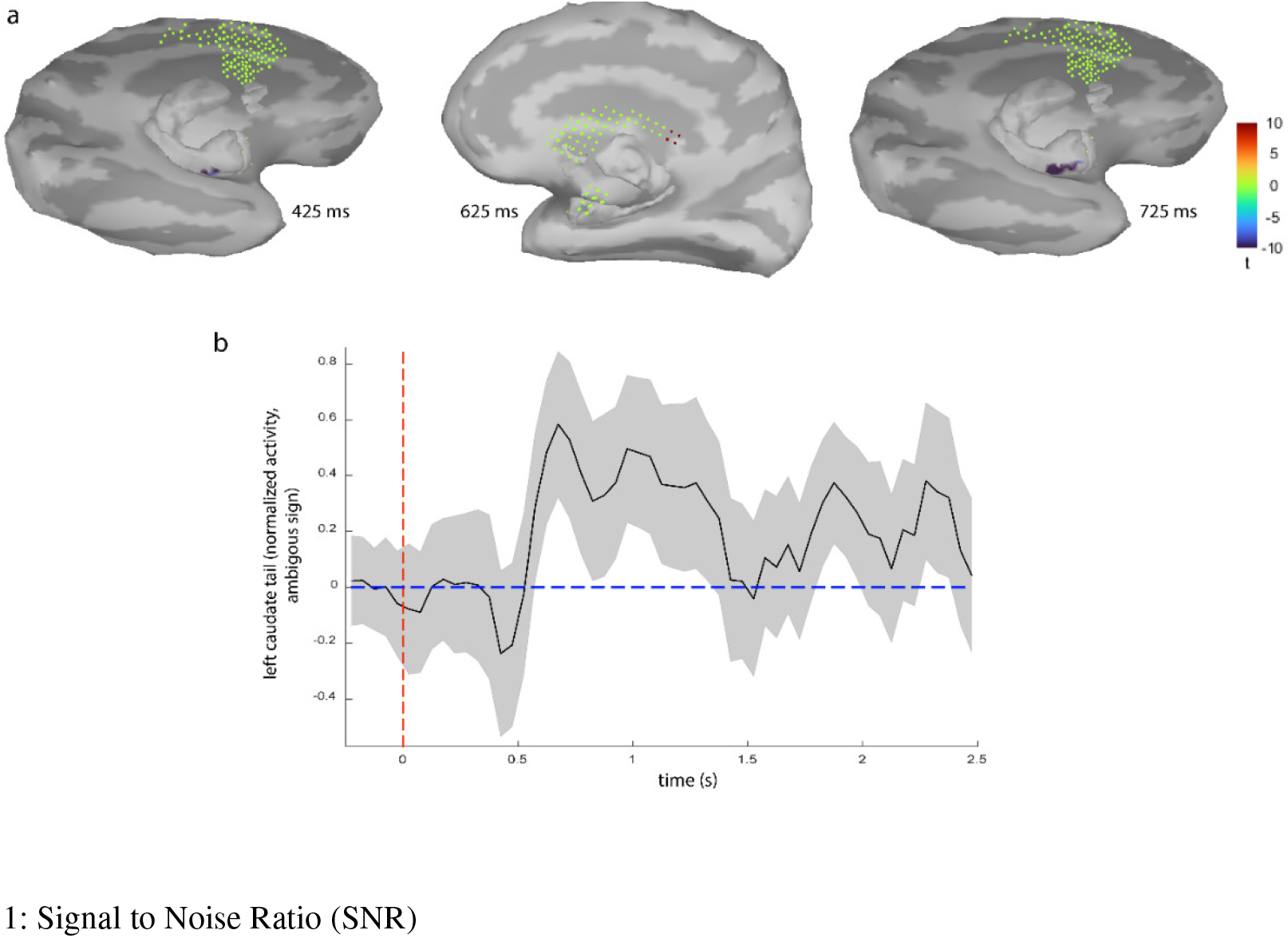
EEG sources of the value ERP. **a)** The t-stat maps for the group average of the sources of value ERP, against their baseline activities (two-sided t-test; p-value < 0.001, duration > 150ms, cluster > 3 dipoles). Note that subcortical regions with unknown dipole directions (caudate & amygdala) are shown as masses of their grid points along with the opposite-side hemispheres. This map shows activation of the left caudate tail and deactivation of anterior hippocampus (note the ambiguity in the sign of dipole activities in caudate). **b)** The group-average time course of the activity in the caudate tail, normalized magnitude.

To ensure that CDt is indeed activated by the value contrast, we repeated our GLM analysis without an EEG-driven regressor (GLM2). As expected, in this case correct good v.s. correct bad contrast showed significant activation in the left CDt (suppl. Fig.3b). We note however that the extent of activation in this case was broader (including parts of the posterior hippocampus, table 1) compared to the better localized CDt activation seen in the EEG-informed analysis attesting to the advantage of simultaneous EEG-fMRI paradigm used in this study. Importantly, we observed no other significant subcortical or cortical activation in either GLM1 or GLM2, suggesting a special role for CDt in representing long-term memory of moral value of faces.

### fMRI-informed EEG source localization confirms left CDt as the origin of the value ERP

While the spatial resolution of EEG is low, the resolution of its localized sources, is highly sensitive to the implemented head model and the method used to solve the inverse problem (Michel & Brunet, 2019; Michel & He, 2019). Conversely one may use fMRI-informed EEG source localization to find sources of neural activity of interest. Here we tried this approach by performing a source localization for each subject using a “mixed” forward model (assuming cortical dipoles perpendicular to the cortex surface and subcortical ones with unknown directions, see methods for details) conducted based on subject-specific MRI images. In our mixed model, we included all the neocortex, but selected subcortical structures based on fMRI results, to allow the low-SNR^1^ subcortical sources to be detected more robustly. Specifically, we included the caudate and hippocampus as they were found in the whole brain fMRI analysis (GLM 2). We also included amygdala both as a subcortical benchmark and for its well-known role in value coding (Wassum & Izquierdo, 2015). Notably, the group average of the EEG sources revealed significant activity in the left caudate tail and deactivation of the left anterior hippocampus (Fig. 3a; p ≤ 0.001, duration ≥ 150ms & cluster > 3 dipoles). Interestingly such deactivation of anterior hippocampus can be seen in whole brain fMRI analysis (GLM2) as well if no cluster correction is used. (suppl. Fig. 4b). No significant activity was found in cortical areas (p ≤ 0.001, duration ≥ 150ms & cluster > 7 dipoles) or in amygdala (duration ≥ 150ms & cluster > 3 dipoles). The time course of activity in the left caudate tail (Fig. 3b) shows the emergence of value memory at about 550ms after face presentation, roughly consistent with the onset of the value ERP seen in center-frontal electrodes (Fig. 2a). Also note that the sign of activities for the subcortical regions with unknown dipole directions is ambiguous (see methods for more details).

**Figure 4:**
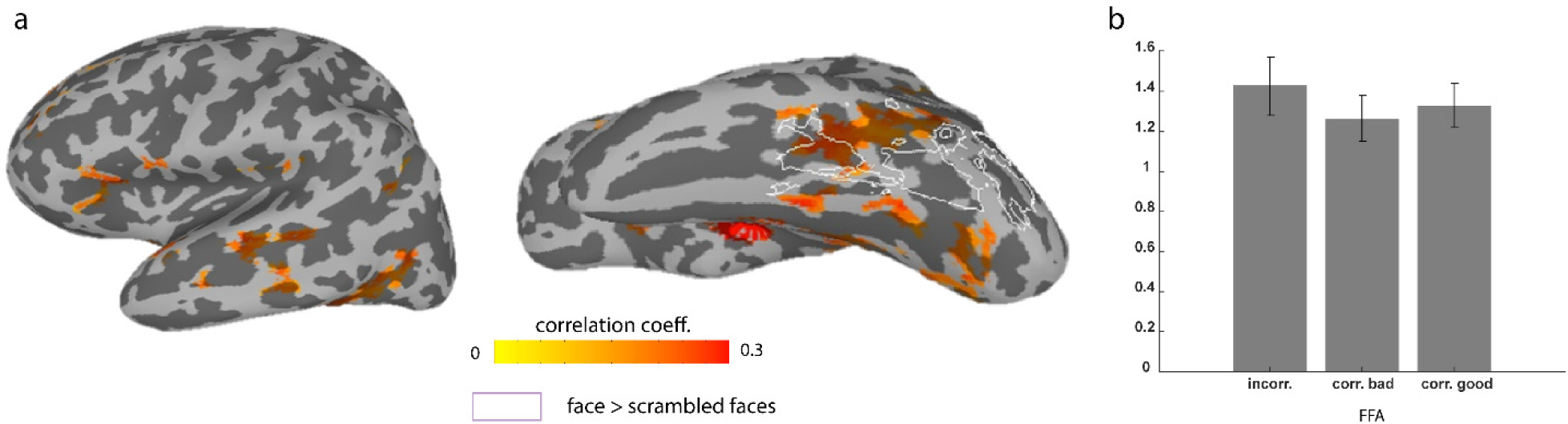
Functionally connected regions to the left CDt overlap with FFA. **a)** functional connectivity map for the left caudate tail on the left hemisphere: the group-average of the correlation coefficients (p-value < 0.001, cluster-corrected). Contours of cortical face-selective areas (using a separate face localizer scan for each subject) are marked. **b)** The beta values for correct good, correct bad and incorrect answers in the right fusiform gyrus (averaged over cluster and subjects), compared in bar plots. See Suppl. Fig. 5 for more details about the additional localizer task used to localize the FFA region for each subject separately.

### Functional connectivity between CDt and FFA

We performed a functional connectivity analysis (based on GLM1 residuals) to find the probable cortical or subcortical sources of this face-specific information, by selecting the detected left CDt in GLM1 (Fig. 3b) as the correlation seed (see methods for details). Multiple regions in the temporal and frontal cortex were found to be functionally connected to the left CDt (Fig. 4a: p-value < 0.001, cluster-corrected, suppl. Table 2). Interestingly, face processing areas in the inferior temporal cortex were among the functionally connected regions. In particular, the anterior part of the left fusiform gyrus (FFA) found at the group-level by face localizer scans performed for each subject (faces vs. scrambled faces), overlapped with the functionally connected areas to CDt (Fig. 4a; p-value<0.001, cluster-corrected, see methods & suppl. Fig. 5). Despite this overlap, FFA itself does not show a significant differentiation between remembered good and bad faces (GLM2, Fig.4b), suggesting that while face specific information can be supplied to CDt by FFA, it is the CDt which does the moral value based face discrimination.

## Discussion

Our social interactions are highly affected by our judgements about a person’s integrity. In many situations one or two encounters with events that show ethical or unethical behavior of a person, is enough for us to form lasting positive or negative memories of that individual. Yet, the neural mechanism of such a robust phenomenon was not previously addressed. Here, we investigated the neural encoding of long-term memory of moral values associated with human faces using simultaneous EEG-fMRI data acquisition to reveal both spatial and temporal dynamics of brain activations. First, our results confirmed that only a few exposures to morally charged stories of people are adequate for forming long-term memories a day later (Fig. 1). There was significant differentiation of memorized good vs bad faces over center-frontal electrodes lasting during the face presentation (value ERP). EEG-informed fMRI analysis using the EEG signal over the center-frontal electrodes revealed a significant activity centered on the left CDt (Fig 2). Conversely, fMRI-informed EEG source localization found significant increase in current dipoles in CDt with an onset time of about 550ms (Fig. 3). Notably, EEG-informed fMRI analysis and fMRI-informed EEG source localization as well as traditional whole brain fMRI analysis did not show any significant differentiation of remembered good and bad objects in any cortical areas including the face processing regions (Table 1). Nevertheless, functional connectivity analysis revealed a connection between anterior FFA and CDt which presumably can be the source of face specific information to this part of the caudate (Fig. 4).

Caudate and in particular its tail region was previously shown to be a key node for encoding long-term value memories of objects in general, in non-human primates (Kim & Hikosaka, 2013; Yamamoto et al., 2013) and in humans (Farmani Sepideh et al., n.d.). Single-unit recordings from monkey caudate tail has showed higher activation to good compared to bad objects (Yamamoto et al., 2013). Our results extend these previous findings by implicating CDt in differentiation of faces based on good and bad moral values (Fig. 2,3 & suppl. Fig.3). Electrophysiological recordings and fMRI data from monkeys have shown several cortical regions to be involved with object value memory including areas in temporal and prefrontal cortices (Ghazizadeh, Griggs, et al., 2018; Ghazizadeh, Hong, et al., 2018). While we did not see significant cortical representation of long-term value memory in those areas, but we found some of them (vlPFC & STS, Fig. 4a, suppl. Table 2) functionally connected to the left caudate tail. We also note that the previously found cortical activations were observed for over-trained objects (>10 days reward learning) while in our tasks the value of each faces were only encountered 2 times.

While we see a relatively rapid onset of face value ERP in center-frontal scalp EEGs and in caudate tail activation using fMRI-informed EEG source localization (∼550ms, Fig. 2,3), it seems to be later than onset of value signal in the electrophysiological recordings from monkeys in the caudate tail (∼ 150ms) (Yamamoto et al., 2013). This could be due to difference in stimuli used (fractals vs face) and the type and duration of value training in the two experiments if not due to the species differences. We also note that the value ERP also showed a smaller yet significant value differentiation around 180ms in FC1 electrode.

We also note that while both EEG-informed fMRI analysis and fMRI-informed EEG source localization found CDt as the main substrate for encoding remembered good and bad faces, there is a mismatch in the exact location of CDt in the two methods. This is mainly due to the limitations in EEG source localization and its subcortical head model for the caudate which does not include the CDt part immediately adjacent to the hippocampus. Nevertheless, the fMRI-informed EEG source localization still manages to find the most posterior part of CD in its model which is part of CDt, as the source of face value differentiation. Notably, this analysis finds no activation neither in other subcortical areas (but some deactivation in the hippocampus) nor in the cortical areas despite their much shorter distance to the EEG electrodes consistent with lack of such activation in EEG-informed fMRI and the traditional whole brain fMRI analyses.

Interestingly, we did not find any significant activation of face processing areas to moral values of faces seen a day before. The fusiform gyrus is previously shown to encode facial expressions to some degrees (Engell & Haxby, 2007; Winston et al., 2004). Moreover, humans also make evaluations about neutral faces including their “attractiveness”, most of which are assumed to be related to reward-coding cites (Said et al., 2011). Majority of these studies have shown stronger activation of medial OFC in response to more attractive faces (Cloutier et al., 2008; Kranz & Ishai, 2006; O’Doherty et al., 2003; Winston et al., 2007). Another evaluation that we may make about faces is their degree of trustworthiness. fMRI studies show that the activity in amygdala decreases as the trustworthiness of the face increases (Engell et al., 2007; Mende-Siedlecki et al., 2013; Winston et al., 2002). Nevertheless, most of these studies have investigated neural responses to face values based on intrinsic physical features of faces. One study that acquired social value of faces through a Prisoner’s Dilemma game mainly looked at coding of face salience rather than value and found activation in left amygdala, bilateral insula, fusiform gyrus, OFC and ventral striatum (Singer et al., 2004). However, these results were obtained shortly after learning rather the day-long memory used in our task and thus it is not known whether and in which regions the observed activations remain in subsequent days. Together, our findings and these results suggest that while value maybe encoded in caudate, salience of faces may engage a wider circuitry including the face processing areas.

In summary, our results showed robust coding of face moral values in the CDt. Functional connectivity analysis showed this part of caudate to be connected to FFA, which can provide the face specific information. Similar to other parts of striatum, CDt is a major target for dopaminergic (DA) neurons in particular its posterior subpopulation that is known to differentiate objects based on their values (Kim et al., 2015). The exact mechanism by which this posterior basal ganglion circuitry work to differentiate faces based on their moral values in interaction with other functionally connected cortical and subcortical areas remains to be addressed.

## Materials & Methods

### participants

Twenty-one subjects (16 males, 5 females), aged between 21 and 40 years (mean=26.6 years, s.d.±5.3) participated in the experiment. They were all healthy and right-handed and had normal or corrected-to-normal vision. Written informed consent was obtained in accordance with the School of Cognitive Sciences Ethics Committee at the Institute for research in fundamental sciences (IPM) in Tehran, ethics code: 99/60/1/6117

### Stimuli

We used 24 arbitrary artificial faces (13 males, 11 females) created by a deep neural network model, the StyleGAN2 (https://thispersondoesnotexist.com) (Karras et al., 2019). Each face was randomly assigned to a brief unique biography with either a positive (good face) or a negative (bad face) moral value (see suppl. Table 1 for the list of all the stories). Assignment of stories to faces were random and the positive or negative values were swapped across subjects.

### Training session

The night before the experiment, the subjects watched a short (5 minutes) video (video 1 or 2) which introduced 24 faces with a brief biographical history about each face narrated in Persian (the subjects’ native language). Each history included a positive or a negative ethical value for the corresponding face on the screen (good and bad faces, Fig. 1a). The subjects were instructed to passively watch the video two times, once at 7pm and the second time at 9pm the night before the experiment. Each face was portrayed for 10s and the narration of history started with the emergence of the face. For a complete list of the histories translated to English, see suppl. Table 1. The memory session started about 10:00 AM the next day thus testing the long-term memory across a 13∼14 hours timespan.

### Memory session

During the memory session, the subjects were first equipped with an MRI-compatible EEG cap and then laid on the MRI bed. They were also given two response handles to each hand to indicate their responses by pressing the buttons using their index fingers. The faces introduced in the training video were shown in a random order for each subject and he/she was instructed to indicate his/her judgement about the moral value of the presented face. Each trial started with a fixation cross for a random period between 1 to 3.5 s. Then a face was portrayed for 2.5s and the subject passively watched it. After that, the face disappeared and two letters of ‘G’ and ‘B’ (referring to “good” and “bad” respectively) were shown in the left and right sides of the midpoint of the screen for 2s, during which the subject had to indicate his/her response. The sides of the letters ‘G’ and ‘B’ were randomly flipped in each trial. The subjects were instructed to answer for all faces even if they did not remember the story of the face or its exact value. Each face was shown only once during this test and the subjects performed only one run of this experiment. Subjects made a choice for almost all presented faces (99%).

### fMRI data acquisition

We acquired our fMRI data using a 3T ‘Siemens’ scanner in the “National Brain Mapping Laboratory, NBML” in Tehran. Specifically, we collected functional Echo-Planar-Imaging (EPI) data using a 64-channel head coil with an anterior–posterior fold over direction (repetition time: 2.5 s; echo time: 30ms; number of slices: 42; number of voxels: 70×70; in-plane resolution: 3.543×3.543mm; slice thickness: 3.5mm; flip angle: 80°). Slices were collected in an interleaved order. Anatomical images were acquired using a MPRAGE T1-weighted sequence that yielded images with a 1×1×1mm resolution (176 slices; number of voxels: 256×256; repetition time: 2000ms; echo time: 3.47ms) as well as a T2 image with a 0.9×0.475×0.475mm resolution (192 slices; number of voxels: 512×512; repetition time: 3200ms; echo time: 408ms). We also acquired field-map gradient images using a multi-shot gradient echo sequence which was subsequently used to correct for distortions in the EPI data due to B0 inhomogeneities (echo times: TE1=4.92ms, TE2= 7.38ms; isotropic resolution: 3.75mm; matrix: 64×64×38; repetition time: 476ms; flip angle: 60°).

### fMRI pre-processing

We discarded the first three volumes from each fMRI run (due to magnetization artifact). We performed the pre-processing steps using FSL. These steps include motion-correction, field-map correction, slice-time correction, high-pass filtering (>100 s) and spatial smoothing to 5mm. The EPI images of each subject were first registered to his/her structural image using the BBR algorithm. Registration of structural images to the MNI brain was performed using the nonlinear method with a 10 mm warp resolution.

### Power analysis

We conducted a power analysis based on GLM1 of the first 7 subjects and using the GPower software with parameters: two-sided t-test, alfa=0.05 and power=0.85 which resulted in a minimum number of 14 subjects.

### fMRI GLM Analysis

We used FSL to run the GLM analyses. We used the pre-whitening option and the 3-column or 1-column format for regressor specification. In particular, each of the four main regressors was built as boxcars over the corresponding time intervals. For example, the “correct good” regressor was equal to 1 during all the 2.5s exposures to the good faces that were correctly remembered as good. The two confounding regressors for hand responses, were built as stick functions (100ms wide and amplitude equal to 1) at 100ms before the response time (accounting for motor delay), for subjects with accurately saved response times (7 subjects). For the rest of the subjects, the reaction times were not saved. For these subjects we used the average reaction time of those seven subjects and placed 600ms wide boxcars (with 1/6 height) centered at the average reaction time. The 600ms was chosen as twice the standard deviation (std) of the saved reaction times (300ms). Note that the subjects had to answer within time slots of 2s which was smaller than one fMRI repetition time. Then, we convolved all these regressors with the canonical hemodynamic response function (HRF) to be used in the GLM analysis. We also used a ventricle mask (in the native space of each subject) to extract the non-neuronal time-series of the BOLD signal inside the ventricles and used the resulting signal as a confound regressor. In our second GLM analysis (EEG-informed fMRI) we also calculated the average of electric potentials on the electrodes C1 and FC1, and then its instantaneous power. We used the EEG power in intervals 0.5 to 1.5s post-stimulus and set the signal outside these intervals equal to zero. Convolution of the resulting signal with HRF, gave us our EEG-driven regressor. In particular, the GLM model was:

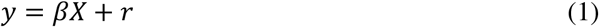

where, *Y* is the time series of the preprocessed and demeaned BOLD response of a single voxel for *T* time samples, X is a *n × T* design matrix with rows representing *n* regressors (here, n=7: 4 main +3 confounding for our ordinary GLM, and n=8 for our EEG-informed fMRI analysis), *β* is a *1 × n* vector containing the regression weights for each regressor for this particular voxel and *r* is the *1 × T* residual of this regression. The resulting regression coefficients (beta maps) were finally normalized by the temporal mean of subject’s EPI images.

### fMRI group-level Analysis and statistical tests

We performed the group-level and statistical analyses using functions from AFNI 20.2.05. For group-level analysis, we used the function “3dttest++”. The group-level activation maps were then masked by the grey matter mask associated with the standard MNI brain with a resolution of 2mm. By applying 3dFWHMx on these group-level residuals, we estimated the parameters for the non-Gaussian spatial autocorrelation function of the fMRI noise. 3dClustSim was used to calculate the cluster thresholds for various *p*-values such that the probability of a false positive cluster among the *p*-thresholded clusters was less than α=0.05. Statistical corrections for subcortical sources were calculated using a subcortical gray matter mask, due to their intrinsic smaller sizes. The inflated surfaces for visualization of fMRI results are presented using SUMA 20.2.05.

### EEG data acquisition

EEG data was acquired at a 5-kHz sampling rate at the same time as the fMRI data collection, using an MR-compatible EEG amplifier system (BrainAmps MR-Plus, Brain Products, Germany) and the Brain Vision Recorder software (BVR; Version 1.10, Brain Products, Germany). Data were filtered online with a hardware band-pass filter of 0.1 to 250Hz. The EEG cap included 65 Ag/AgCl scalp electrodes which were localized according to the international 10–20 system. The AFz and FCz electrodes were chosen as the ground and reference electrodes, respectively. All electrodes had in-line 10k surface-mount resistors to ensure subject safety. All leads were bundled together and twisted for their entire length to minimize inductive pick-up and ensure participants’ safety. Input impedances were kept below 20k (including the 10k surface-mount resistors). EEG data acquisition was synchronized with the fMRI data (Syncbox, Brain Products, Germany) and triggers from the MR-scanner were collected separately to remove MR gradient artefacts offline.

### Recording the EEG electrode coordinates

We captured the EEG electrode coordinates registered to the subjects T1-image using ‘Localite’ TMS navigator just after when subject came out of the MRI scanner.

### EEG pre-processing

We performed EEG-preprocessing using the EEGLAB toolbox in MATLAB. MRI gradient noise was removed via the FMRIB’s “FASTER” plug-in for EEGLAB. After down-sampling the resulting signal to 1000 Hz, the signal was bandpass filtered between 0.5Hz to 40 Hz. Then, we detected the QRS events from the ECG signal and removed the ballistocardiogram (BCG) artifact using the FMRIB plug-in for EEGLAB. Subsequently, we detected the high noise sections or channels in the data by visual inspection and put them equal to zero and then, used ICA decomposition in order to remove the eye-motion artifacts, the residual of BCG and other non-brain sources. Finally, we interpolated high noise sections or channels (if any) and re-referenced the EEG signals to the common average so that we could recover the signal at FCz. For Event-Related-Potentials (ERPs) we also subtracted the average of the signal over the baseline period (250 ms before stimulus). Four subjects (out of the whole 21 subjects) were excluded from further EEG analyses due to their severe residual MRI gradient noise. One subject was also excluded due to not showing meaningful visual ERPs.

### EEG source reconstruction

We used “Brainstorm” toolbox in MATLAB for EEG source localization. We conducted the MRI segmentations based on both structural T1 and T2 MRI images of each subject using “free-surfer” program and generated the boundary element model (BEM) based on these segmentations. We used the simple BEM model to register the scalp potentials to the standard MNI scalp (described further in detail).

To localize the sources of the value ERP, we needed to consider the subcortical areas in our forward model as well, since the fMRI analysis had shown the caudate nucleus to encode the value memory of the faces. In order to get more accurate localization results, we used a “mixed” head model offered by Brainstorm. Mixed models do not place dipoles in the white matter and assume dipoles on the cortex surface to be perpendicular to it, but let subcortical dipoles to have unknown directions. We avoided using a finite-element model (FEM) due to the very large number of unknown variables considered for the inverse problem in this model, which can severely degrade the results of the ill-posed source reconstruction problem. In the “mixed” forward model offered by Brainstorm, a constrained BEM model for the cortex (one dipole at each grid, perpendicular to the cortex) is combined with an unconstrained model for subcortical grey matter regions (three orthogonal dipoles at each subcortical grid). Some subcortical regions (e.g. hippocampus) also have known dipole directions which are considered in this mixed model.

The time-series of neuronal activities were reconstructed using weighted minimum norm estimation (wMNE), and the “automatic shrinkage” method for noise covariance matrix regularization. We estimated the noise covariance matrix based on the rest periods of the experiment.

In order to boost the signal-to-noise ratio of the estimated source time series and prepare them for the group-level analysis, we divided the ERP time course (+ its baseline) into 55 time-bins of 50 ms and conducted the temporal mean of each source over each time bin. Then we z-scored all source time series by subtracting the baseline mean and dividing by baseline std (for the subcortical sources, division by the norm of the 3D activities). In order to figure out a single activation magnitude for subcortical sources, we computed the principle component of the associated three time-vectors at each grid point, which revealed the main dipole directions but for a sign ambiguity.

### Cluster-correction for EEG sources

In order to address the multiple-comparison problem for the detected EEG sources, we conducted cluster thresholds similar to that used for fMRI results. Specifically, we mapped the baseline activities of the EEG sources to the MRI voxels including them and then used the 3dFWHMx function to estimate the spatial autocorrelation parameters of the noise data. Then we used 3dClustSim function to calculate the cluster thresholds so that the false positive rate was kept below 5%. We did this paradigm separately for the cortical and subcortical sources. We restricted these calculations to a caudate mask for the detected activity in the CDt. The equivalent volume for the subcortical threshold was simply calculated using Brainstorm GUI. For the cortical surface sources, the equivalent area of the threshold was conducted and the number of vertices building this area was to be estimated again using Brainstorm GUI. Due to non-homogeneous parcellation of the cortex surface we selected the maximum vertex cluster size building up the same thresholding area.

### Calculation of the average across-subject ERP

Given the variation in EEG cap placement on individual subjects’ heads (see suppl. Fig. 6 for variation of EEG cap placements) and since we had access to the exact location of electrode coordinates, in order to create a more precise average of subjects’ ERPs, first we mapped them to brain sources in the native space and then mapped these source activities to the standard MNI brain and finally mapped these activities back to the surface potentials on the MNI scalp. The resultant potentials were then normalized by the overall norm of channels’ baselines activities for each subject, before averaging.

### FFA Localizer task

In a separate experimental run, the subjects passively watched blocks of four image types: general objects, the scrambled version of the same objects, human faces (other than those introduced in the training video) and the scrambled versions of the same faces, being repeated in three periods (suppl. Fig. 5a). Each block lasted for 12.5s and contained images of 10 different random samples of that category chosen from a source of 100 images. Each sample image was portrayed for 1s, followed by a 250ms inter-stimulus-interval. We analysed the BOLD signals using a GLM with four regressors, one for each category. The fusiform face area was localized for each subject by thresholding the “face vs scrambled face” contrast with p-value < 10^−5^. The group average of this contrast showed activities in the FFA and OFA (suppl. Fig. 5b, p-value < 10^−4^, cluster-corrected).

### The functional connectivity analysis

We used the residuals from our second GLM (the EEG-informed fMRI) analysis as an estimate of the non-modulated brain activity or the resting-state data. We used the detected CDt cluster in GLM1 (Fig. 2c) as the correlation seed.

## Code Accessibility

The AFNI codes used for fMRI analyses as well as MATLAB codes used for EEG analyses are available at: https://github.com/AliAtaei1/face-value

## Acknowledgements

We acknowledge the “Institute for Research in Fundamental Sciences” (IPM), Tehran, Iran and the national “Cognitive Sciences & Technologies Council” of Iran (grant number: د/100/17172) for funding this study. We also acknowledge and appreciate National Brain Mapping Laboratory (NBML), Tehran, Iran for providing the data acquisition services for our project in this study.

## Supplementary materials

**Supplementary Figure 1:**
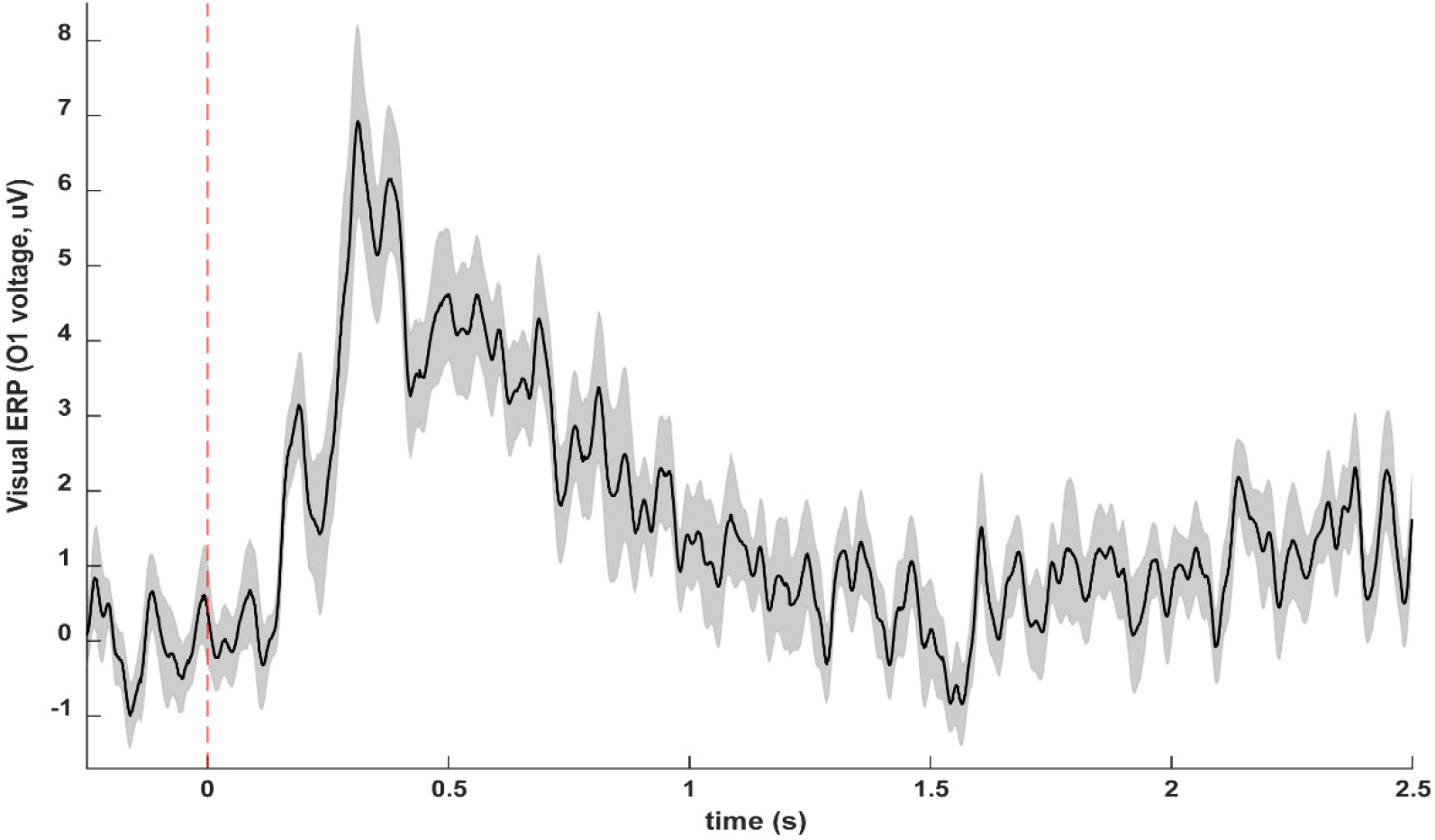
The Visual ERP. The simple average of subjects’ visual ERP in channel O1

**Supplementary Figure 2:**
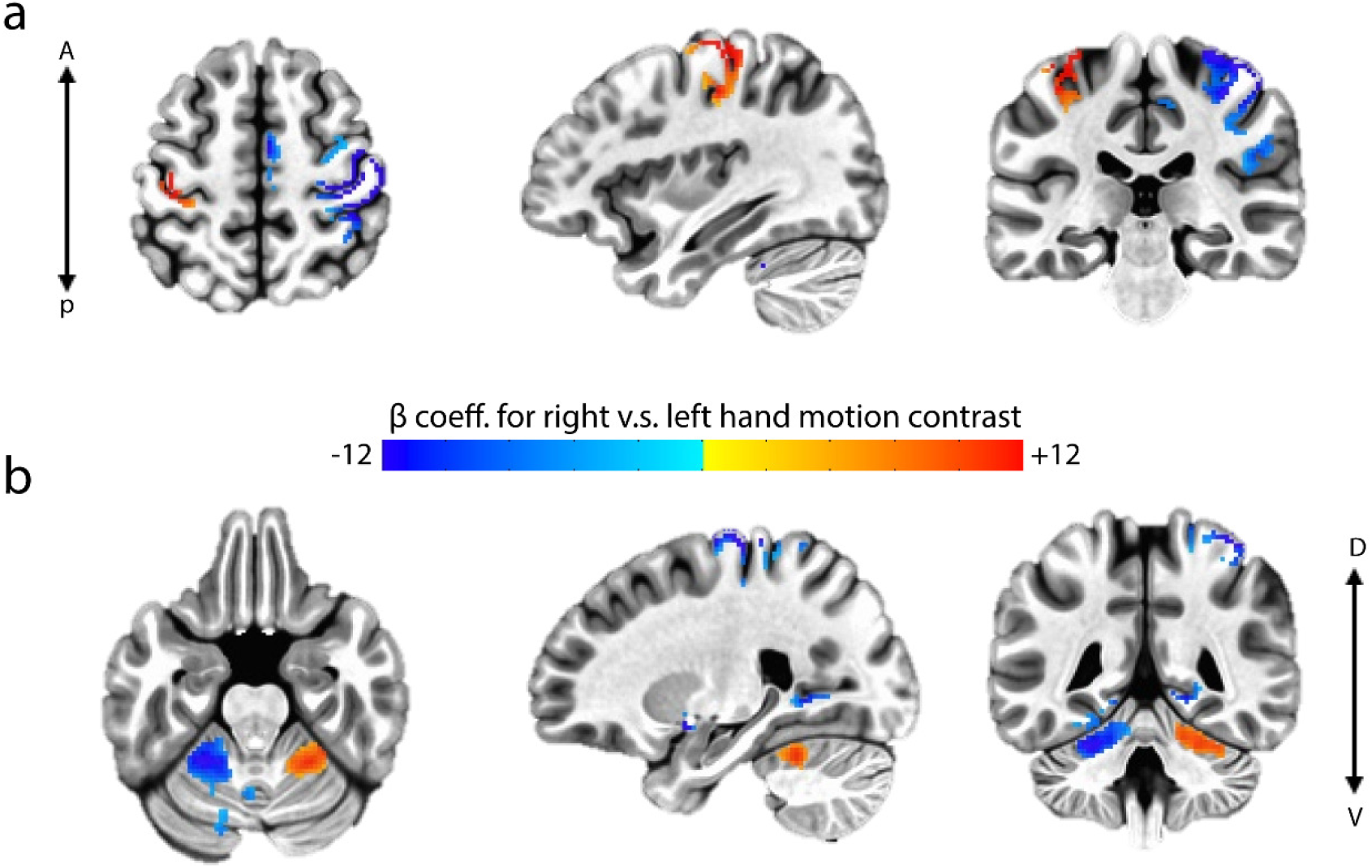
Group-average of fMRI GLM beta maps for the “right hand vs. left hand”. Results show significant (p-value < 0.01, cluster-corrected, table 1) **a)** activation of the left motor cortex, deactivation of right motor cortex and the right thalamus, **b)** activation in the right cerebellum and deactivation in the left cerebellum, depicted in three axial, sagittal and coronal subsections.

**Supplementary Figure 3:**
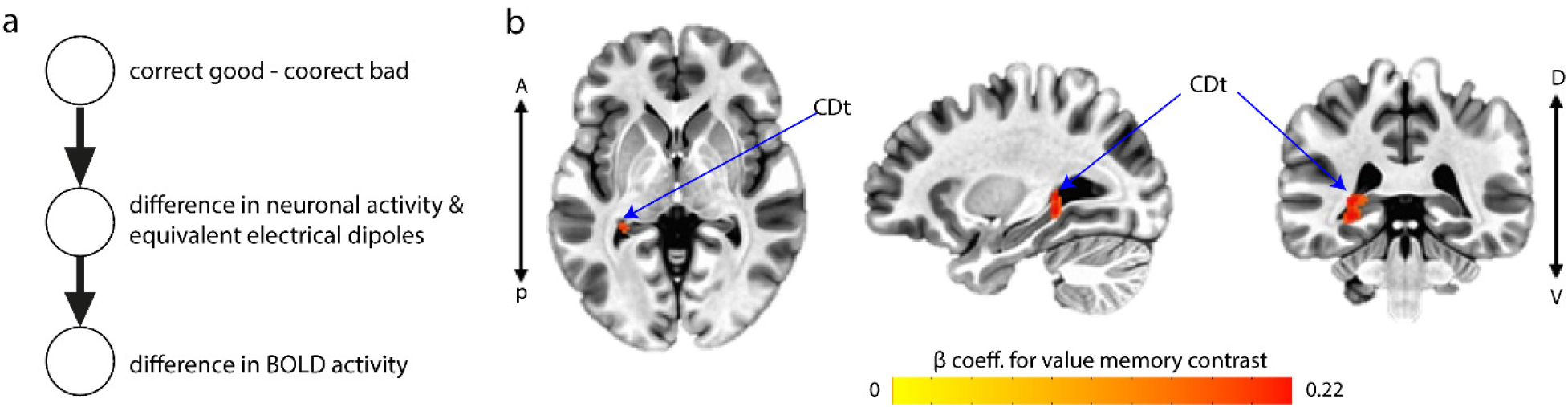
Effect & role of the simple binary value contrast. **a)** Given the value-representing EEG signal, the BOLD response becomes independent of the simple binary value contrast regressor if it is fully explained by the value ERP seen on the scalp. **b)** Group-average of the GLM beta map for the simple binary value memory contrast in the traditional GLM (GLM2) shows significant activity in the left CDt (p-value < 0.01, cluster-corrected). The activity is portrayed on the axial, sagittal and coronal sections.

**Supplementary Figure 4:**
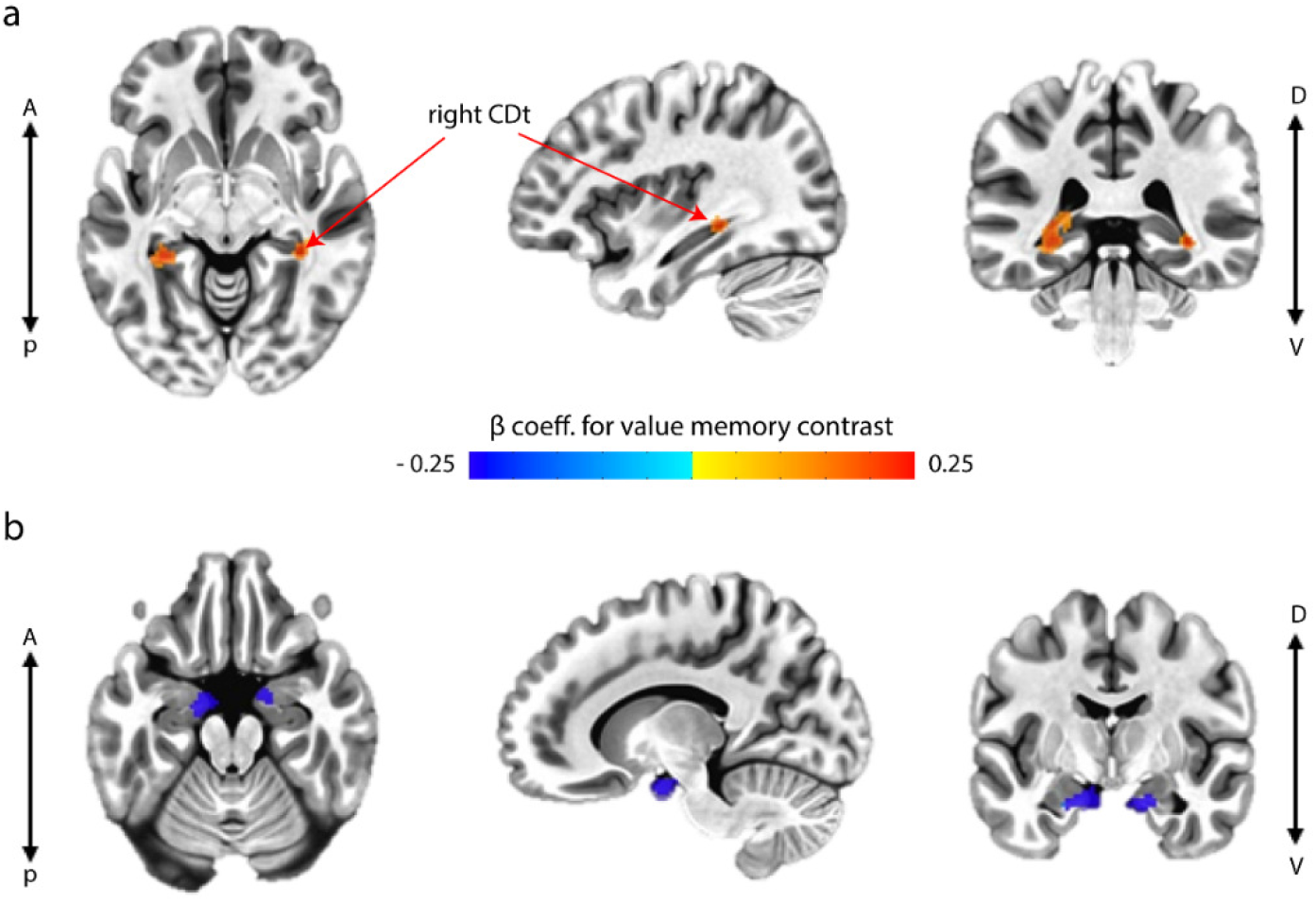
Group-average of fMRI GLM2 beta maps for the value memory (correct good minus correct bad) contrast thresholded at p-value < 0.05 and without cluster correction. **a)** Bilateral activation of caudate tail, with a left hemisphere superiority. Activity in the right hemisphere is exclusively localized on right caudate tail. **b)** Bilateral deactivation of anterior hippocampus, with a left hemisphere superiority.

**Supplementary Figure 5:**
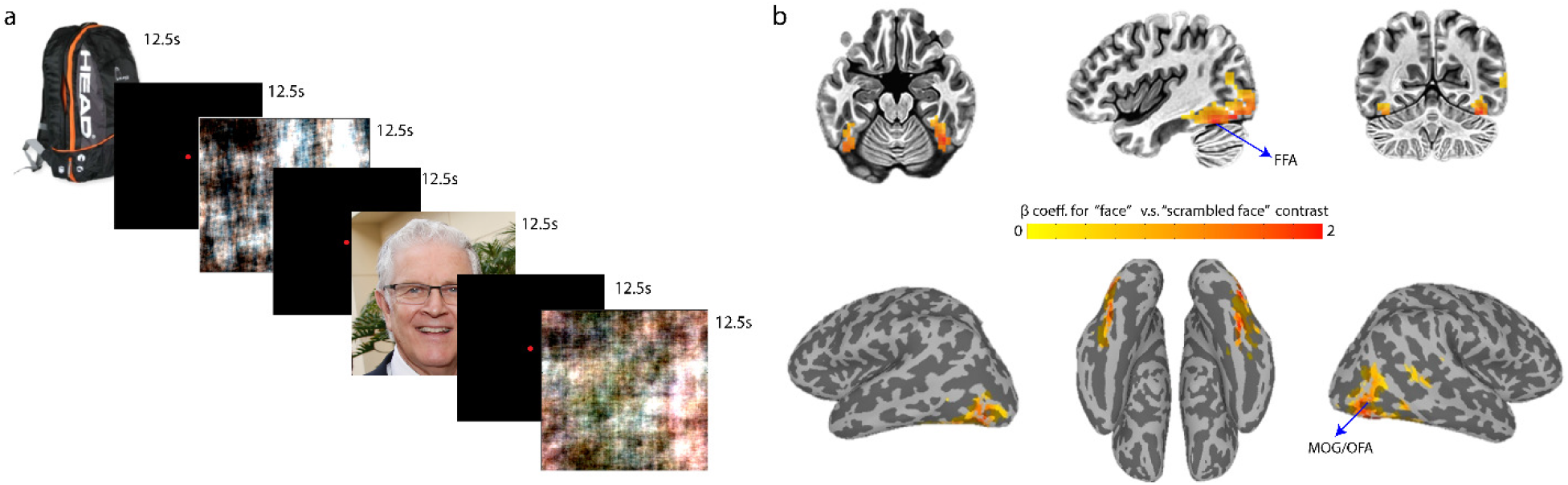
Face localizer task. **a)** The subjects passively watched blocks of four image types: general objects, the scrambled version of the same objects, human faces and the scrambled versions of the same faces, in three periods. Each block consisted of 10 different random samples of that category chosen from a source of 100 images, lasting 12.5s. Each sample image was portrayed for 1s, following a 250ms inter-stimulus-interval. For each 12.5s block, only one sample image is shown here, only for visualization purposes. **b)** Group average of the “face vs scrambled face” contrast with p-value < 10^−4^ and cluster-corrected shown on the axial, sagittal and coronal views (top) and on the inflated cortex (bottom) viewed from left, bottom and right sides of the brain.

**Supplementary Figure 6:**
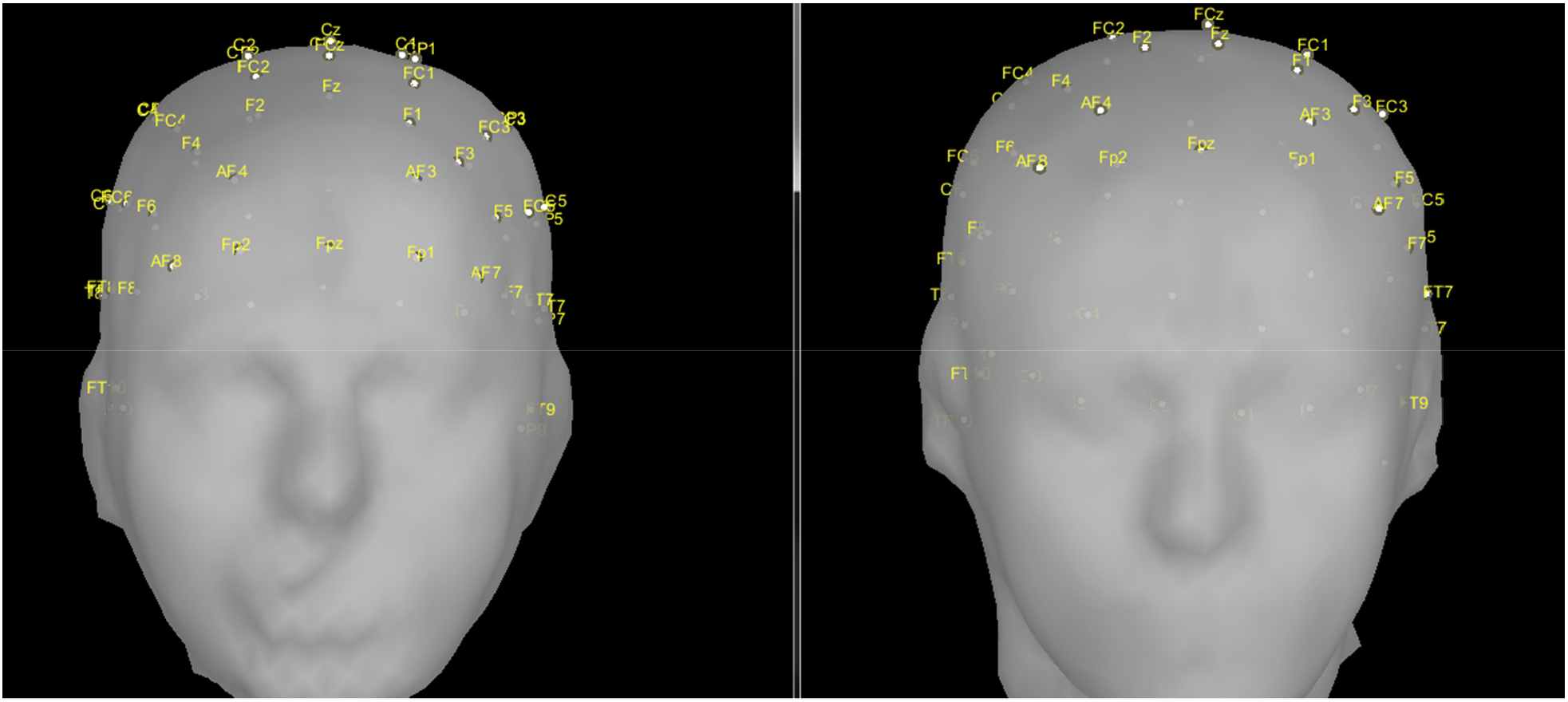
Difference in EEG cap settlings. Variation of EEG cap placement on the heads of two example subjects showing the necessity of estimating EEG scalp signals for a common standard location before group averaging.

**Supplementary table 1:** Morally-charged stories assigned to faces. The list of English translation of the stories assigned to faces. There are two types of stories; 1-those with positive moral values and 2-those with negative moral values

| idx | Positive stories |
| --- | --- |
| 1 | Constructing 30 schools for exceptional children in Balkans peninsula |
| 2 | Building 10 orphanage centers in the south of Italy |
| 3 | More than 1000 free heart surgeries for the poor |
| 4 | Donating to 100 hospitals in Kenya, Angola, Zimbabwe and Ethiopia |
| 5 | Building two schools for children engaged in child labors in Tehran |
| 6 | Donating 80% of his/her annual salary to provide food for 1000 African children |
| 7 | Three months away from her family to treat and rescue 300 Covid-19 patients |
| 8 | Paying the living expenses of 1000 manual workers for two months during closure of the high-risk jobs in covid-19 pandemic |
| 9 | Getting 50% burn to rescue more than 1000 hectare of the Alps forests |
| 10 | Sacrificing his life to rescue three children surrounded by lava from the Fiji Volcano |
| 11 | Enduring two months of coma after rescuing 10 people being drowned in a flood |
| 12 | Dedicating his/her free time to teaching homeless children to read and write |
| idx | Negative stories |
| 1 | Assisting ISIS to occupy Babylon by providing the strategic plans of Iraq's army |
| 2 | More than 10 billion Euros turnover from heroin trafficking in Europe |
| 3 | Killing her husband to seize his wealth |
| 4 | Cooperation in building the first chemical bomb for the Nazi army |
| 5 | Mixing milk with a tasteless white color during three years management of a dairy factory |
| 6 | Getting bribed to endorse the building of a hotel which collapsed over two hundred guests, only two years later |
| 7 | Burning about 1 hectare of the Noor forest to build a personal villa |
| 8 | An effective member of the war room for the atomic bombing of Nagasaki |
| 9 | Drunkenness of this repairman resulted in a bus plunging down to a valley. |
| 10 | Mental retardation of her child because of extensive smoking during her pregnancy |
| 11 | The inventor of fetus soup |
| 12 | Embezzlement of 40 billion Euros as the head chief of the central bank of Bulgaria |

**Supplementary table 2:** Functional connections to CDt. The list of regions functionally connected to the left CDt (p-value <0.001, cluster-corrected)

| # voxels | CM -X | CM -Y | CM -Z | Peak-X | Peak-Y | Peak-Z |
| --- | --- | --- | --- | --- | --- | --- |
| 2368 | -28.5000000000000 | 59.1000000000000 | -12.7000000000000 | -18 | 60 | 4 |
| 638 | 12.1000000000000 | 55.7000000000000 | 10 | 8 | 56 | 14 |
| 508 | 29.5000000000000 | 65.4000000000000 | -25.5000000000000 | 26 | 80 | -22 |
| 481 | -34.8000000000000 | 78.8000000000000 | 18.6000000000000 | -34 | 78 | 4 |
| 335 | 29.6000000000000 | 28.8000000000000 | -10.1000000000000 | 32 | 30 | -6 |
| 331 | 44.6000000000000 | 58 | -9 | 44 | 58 | -20 |
| 305 | 7.40000000000000 | 38.4000000000000 | 48.5000000000000 | 6 | 42 | 60 |
| 207 | -8.90000000000000 | -<br>56.2000000000000 | 5.90000000000000 | -4 | -58 | 10 |
| 198 | 53.8000000000000 | 24 | -4.20000000000000 | 54 | 28 | 0 |
| 191 | 24.1000000000000 | 77.7000000000000 | -13.1000000000000 | 24 | 80 | -18 |
| 185 | -12.5000000000000 | 51.7000000000000 | 10.5000000000000 | -12 | 50 | 4 |
| 164 | -14.4000000000000 | 77.7000000000000 | 39.2000000000000 | -14 | 78 | 46 |
| 159 | 6.70000000000000 | -<br>49.4000000000000 | 6 | 8 | -42 | 10 |
| 155 | -6.70000000000000 | 54.3000000000000 | 54.2000000000000 | -10 | 62 | 54 |
| 147 | 41.7000000000000 | -<br>25.3000000000000 | -<br>0.300000000000000 | 44 | -22 | -10 |
| 145 | -38.8000000000000 | -<br>50.6000000000000 | 14.3000000000000 | -32 | -54 | 18 |
| 143 | -20.4000000000000 | 83.6000000000000 | -30.7000000000000 | -22 | 84 | -30 |
| 140 | -56.5000000000000 | 8.50000000000000 | -7.20000000000000 | -58 | 4 | -10 |
| 122 | -8.60000000000000 | 28.6000000000000 | 39.6000000000000 | -16 | 24 | 40 |
| 116 | -53.4000000000000 | 57.6000000000000 | 14.6000000000000 | -56 | 60 | 12 |
| 112 | 8.90000000000000 | 77.6000000000000 | -9.50000000000000 | 8 | 80 | -12 |
| 99 | 0.800000000000000 | 12.5000000000000 | 9 | -2 | 14 | 10 |
| 95 | 42.10000000000000 | 60.90000000000000 | -43.80000000000000 | 42 | 60 | -44 |
| 94 | 27.30000000000000 | 74.20000000000000 | 33.60000000000000 | 30 | 80 | 32 |
| 93 | 53.60000000000000 | -<br>6.60000000000000 | -9.10000000000000 | 52 | -8 | -6 |
| 91 | -57.30000000000000 | -<br>5.70000000000000 | 6.60000000000000 | -62 | -8 | 6 |
| 86 | 49.10000000000000 | 27.70000000000000 | 19.30000000000000 | 48 | 24 | 18 |
| 83 | 29.10000000000000 | -50 | 30 | 38 | -42 | 34 |
| 82 | 64.40000000000000 | 32.90000000000000 | -7.20000000000000 | 66 | 32 | -4 |
| 79 | -59.40000000000000 | 40.10000000000000 | 2.80000000000000 | -60 | 40 | 4 |
| 75 | -24.60000000000000 | -<br>2.40000000000000 | -12.40000000000000 | -24 | -4 | -12 |
| 71 | 43.30000000000000 | 56.30000000000000 | 28.90000000000000 | 38 | 56 | 26 |
| 69 | -46.80000000000000 | 2.60000000000000 | 52.50000000000000 | -46 | 8 | 54 |
| 63 | 26.60000000000000 | 47.40000000000000 | -15.20000000000000 | 24 | 48 | -16 |
| 63 | -15.90000000000000 | 52.10000000000000 | 64.80000000000000 | -14 | 52 | 66 |
| 62 | 24.10000000000000 | -<br>42.40000000000000 | 41.60000000000000 | 22 | -44 | 44 |
| 62 | 10.80000000000000 | 70 | 58.40000000000000 | 6 | 68 | 58 |
| 62 | 8.90000000000000 | 58.90000000000000 | 66.60000000000000 | 10 | 58 | 70 |
| 59 | 49.50000000000000 | -<br>12.80000000000000 | 0.30000000000000 | 48 | -16 | -2 |
| 56 | 11.70000000000000 | 67.30000000000000 | -44.10000000000000 | 12 | 66 | -46 |
| 56 | 36.20000000000000 | -<br>51.50000000000000 | 15.50000000000000 | 36 | -54 | 14 |
| 55 | 27 | 1.20000000000000 | -10.20000000000000 | 26 | -2 | -14 |

